# Improve Concentration of Frequency and Time (Conceft) by Novel Complex Spherical Designs

**DOI:** 10.1101/2020.11.23.394007

**Authors:** Matt Sourisseau, Yu Guang Wang, Robert S. Womersley, Hau-Tieng Wu, Wei-Hsuan Yu

**Affiliations:** Department of Mathematics, University of Toronto, Toronto, ON, Canada *Email address:*; School of Mathematics and Statistics, University of New South Wales, Sydney, NSW, 2052, Australia; ICERM, Brown University, Providence, RI, USA *Email address:*; Departments of Mathematics and Department of Statistical Science, Duke University, Durham, NC, USA; Mathematics Division, National Center for Theoretical Sciences, Taipei, Taiwan. *Email address:*; Department of Mathematics, National Central University, Taoyuan, Taiwan; ICERM, Brown University, Providence, RI, USA *Email address:*

**Keywords:** ConceFT, multitaper, quasi-uniform spherical design, synchrosqueezing transform, time-frequency analysis

## Abstract

Concentration of frequency and time (ConceFT) is a generalized multitaper algorithm introduced to analyze complicated non-stationary time series. To avoid the randomness in the original ConceFT algorithm, we apply the novel complex spherical design technique to standardize ConceFT, which we coin *CQU-ConceFT.* The proposed CQU-ConceFT is applied to visualize the spindle structure in the electroencephalogram signal during the N2 sleep stage and other physiological time series.

## 1. Introduction

Quantifying *oscillatory* time series has attracted much attention recently, particularly those having multiple deterministic oscillatory signals with time-varying frequencies and amplitudes and contaminated by noise; for example, peripheral venous pressure (PVP) signal [13], photoplethysmogram [11] and many others. This kind of problem is a generalization of the widely considered *seasonality* problem [1] in statistical society. There have been extensive studies in the time-frequency (TF) analysis community aiming for this kind of time series [7]. In this report, we propose to combine the recently developed novel complex spherical designs and a nonlinear-type TF analysis to stably analyze a noisy time series. We focus on the STFT-based synchrosqueezing transform (SST) as an example of nonlinear-type TF analysis to illustrate the idea, although we can consider other nonlinear-type TF analyses [7]. It is shown in [2] that SST is robust to different types of noise, such as non-stationary and heteroscedastic noises. However, when the signal-to-noise ratio (SNR, defined as 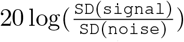, where `SD` means the standard deviation) is low (e.g., below 1 dB), SST may fail. A natural idea to handle time series with a low SNR is taking the *multi-tapering (MT)* technique into account [10, 17]. However, due to the Nyquist rate limitation in the TF representation (TFR) [3], in practice we can only find limited orthonormal windows (i.e. 6 to 10) that have reasonably concentrated supports in the TF domain. Hence, the noise suppression effect by the MT scheme is usually limited. In 2016, driven by the demand to analyze physiological signals with a low SNR, a *generalized MT* was proposed in [4] as a new algorithm, called *concentration of frequency and time (ConceFT).* The basic idea beyond ConceFT is a marriage of the nonlinearity of the chosen nonlinear-type TF analysis and MT. In the generalized MT, we consider *J* orthonormal windows, *h*_1_,…,*h_J_*, where *J* ∈ ℕ, and *randomly* linearly combine them into a new normalized window. By viewing the coefficients of a linear combination of *J* orthonormal windows as a point on the (*J* – 1)-dim real sphere 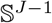, one can obtain as many windows as possible by randomly sampling points 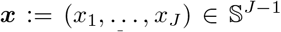 uniformly. In other words, for each ***x***, we have a new window 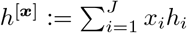. Finally, we average all TFR’s by SST with those windows. While the ConceFT has been applied to a variety of medical signal processing problems [9, 15, 16], for the clinical application purpose, there is still a gap. In fact, due the step of *random* samples from 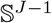, the analysis result varies from one experiment to another and leads to an un-reproducibility issue unless a large number of sampling points are considered. However, this might significantly increase computational complexity. Second, the performance of ConceFT is affected by the approximation quality of numerical integration over the chosen spherical point set. A natural question is then to find a *deterministic* set of points that distributed on 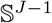 as “uniformly” as possible, and meanwhile have an excellent performance for numerical integration. Below, we consider a novel *spherical design* (SD) [6] scheme on complex spheres [12] to the generalized multitaper scheme to resolve the above two issues.

## 2. Concentration of frequency and time – old and new

For a properly defined function *f*, e.g., a tempered distribution, and a proper window *h*, denote SST of *f* to be 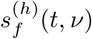, where *t* ∈ ℝ is the time, *v* ∈ ℝ^+^ is the frequency. Fix *N* ∈ ℕ. Take 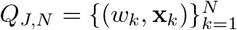 to be a quadrature rule for numerical integration on 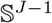 or complex sphere Ω^*J*^ with *N* pairs of weights *w_k_* ∈ ℝ and nodes 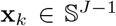 or x_*k*_ ∈ Ω^*J*^.

### Definition 2.1.

For *q* > 0, the ConceFT for the *q*-normed SST associated with *Q_J,N_* for a given proper function *f* is

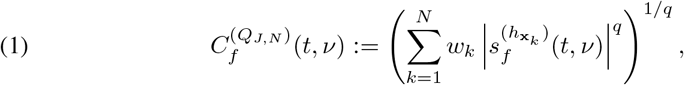

where *t* ∈ ℝ and *v* > 0. We also define the *ideal ConceFT* as

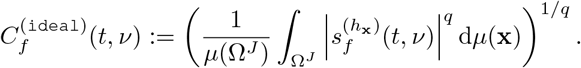

In [4], *q* =1 and *Q_J,N_* is chosen so that {**x**_*k*_} is the set of *N* random uniform (RU) points from 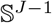 and *w_k_* = 1/*N*. We call this approach the *RU-ConceFT.* Note that in [4], the RU-ConceFT is defined only on *real spheres*. However, in practice we can take the set of N RU points from Ω^*J*^, and we call the resulting algorithm *CRU-ConceFT.* In the report, we propose to take {**x**_*k*_} to be the nodes of a quasi-uniform spherical t design (QU-t-SD) on a real sphere to replace the random sample scheme in [4]. The new algorithm is called *QU-ConceFT.* If we consider a recently proposed quasi-uniform spherical *(k, l)* design (QU-kl-SD) on a complex sphere [12], the algorithm is called *CQU-ConceFT.* When the quadrature rule in Definition 2.1 is a SD, the weights *w_k_* are equal weights 1/*N*. Note that by high precision for numerical integration of QU-t-SD or QU-kl-SD, (C)QU-ConceFT better approximates the ideal ConceFT than (C)RU-ConceFT proposed in [4]. See Appendix for quantitative results on simulated signals, which shows the benefit of CQU-ConceFT and its improvement.

## 3. Numerical experiments

In this section, we show numerical experiments of analyzing simulated signals with noise.

### 3.1. Simulated signals with noise

We use the smoothened Brownian motion as the true signal. This is the model considered in [4] to quantitatively evaluate different algorithms. The signal takes the form

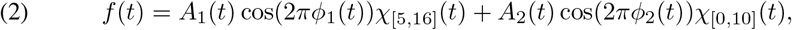

where *t* ∈ [0,16] and *A*_1_(*t*), *A*_2_(*t*), *ϕ*_1_(*t*) and *ϕ*_2_(*t*) are constructed from the same procedure shown in [4, Section 4a].

Specifically, if *W* is the standard Brownian motion defined on [0, ∞), the *smoothened Brownian motion with bandwidth B >* 0 is Φ_*B*_:= *W * K_B_*, where *K_B_* is the Gaussian function with the standard deviation (SD) *B* > 0 and * denotes the convolution operator. Given *T* > 0 and parameters *ζ*_1_,…, *ζ*_6_ > 0, define the following family of random processes on [0, *T*]:

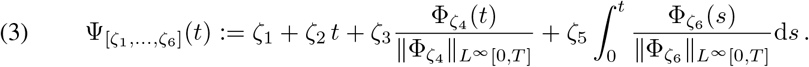

For the amplitudes *A*_1_(*t*) and *A*_2_ (*t*), we set *ζ*_2_ = *ζ*_5_ = 0 and they are independent realizations of ψ_[2,0,1,200,0,0]_ (*t*). To simulate phase functions *ϕ*_1_(*t*) and *ϕ*_2_(*t*), we set *ζ*_1_ = *ζ*_3_ =0 and ψ_[0,*ζ*_2_,0,0,*ζ*_5_,*ζ*_6_]_(*t*) is then a monotonically increasing process. In the examples below, we take *ϕ*_1_(*t*) as a realization of ψ_[0,10,0,0,6,400]_(*t*), and *ϕ*_2_ (*t*) as a realization of ψ_[0,2π,0,0,2,300]_(*t*). The noisy signal is the superposition *Y*(*t*) = *f* (*t*) + *ξ*(*t*), where the noise *ξ*(*t*) is a white mean-zero Gaussian random process with the variance *σ*^2^. The SNR for *Y* is 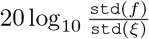, where `std`(*f*) is the standard deviation of *f*. We set the SNR to be 0.11 dB, and realize the stochastic signal *Y*(*t*) for *t* ∈ [1/*ω*_0_,16] with the sampling rate *ω*_0_ = 100 (Hz). Figure 1 shows a realization of the clean signal *f* (*t*), Gaussian noise *ξ*(*t*), and their superposition *Y* (*t*).

**Figure 1.**
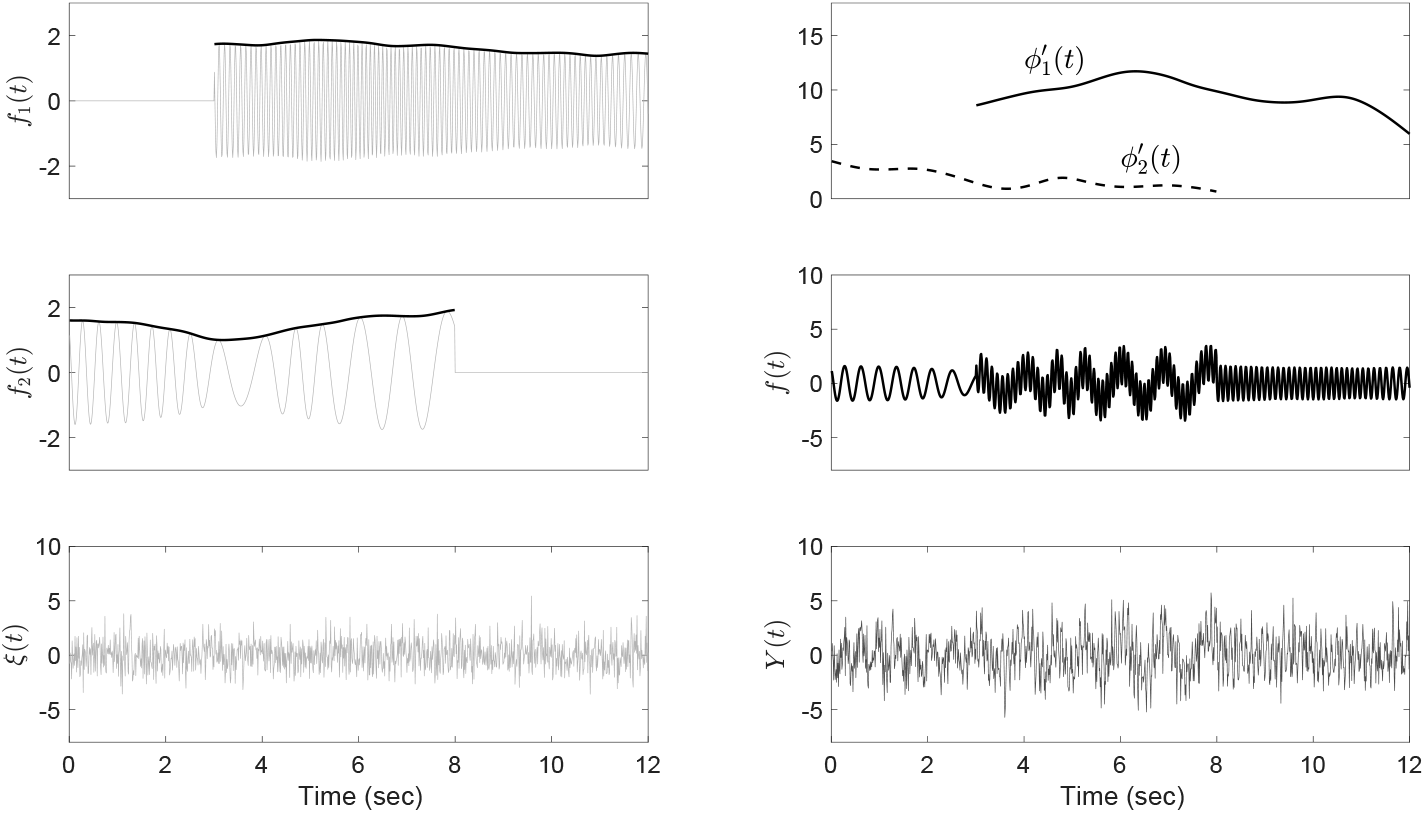
A realization of the stochastic signal *Y*(*t*). In the left column, the first and second subplots are 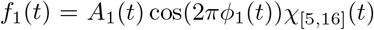 and 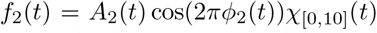 in (2) with *A*_1_ (*t*) and *A*_2_ (*t*) superimposed as black curves, and the third subplot shows a realization of the Gaussian noise *ξ*(*t*). In the right column, the first subplot shows 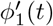 and 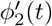, the second subplot shows the clean signal *f* (*t*), and the third subplot is the noisy signal *Y*(*t*) that is the superposition of *f* (*t*) and *ξ*(*t*).

### 3.2. Time-frequency representation

Figure 2 shows MT-SST, QU-ConceFT, CQU-ConceFT [14]^1^, RU-ConceFT and CRU-ConceFT for the realization of *Y* in Figure 1. For MT-SST, we take the first 6 Hermite windows that are the most concentrated in the TF domain among others [3]; that is, we take *J* = 6 tapers. For QU-t-SD, we take a 7-design with 32 nodes on 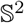. For QU-kl-SD, we take a triangle complex 4-design with 40 nodes on Ω^3^ coming from a real sphere 4-design on 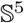. For RU and CRU sets, they are 32 points on 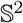 and Ω^3^ respectively. in other words, for RU-conceFT, QU-conceFT, cRU-conceFT and CQU-ConceFT, we take *J* = 3 orthonormal windows. Again, we take the first 3 Hermite windows due to its concentration property in the TF domain. We take *q* = 1 for all cases.

**Figure 2.**
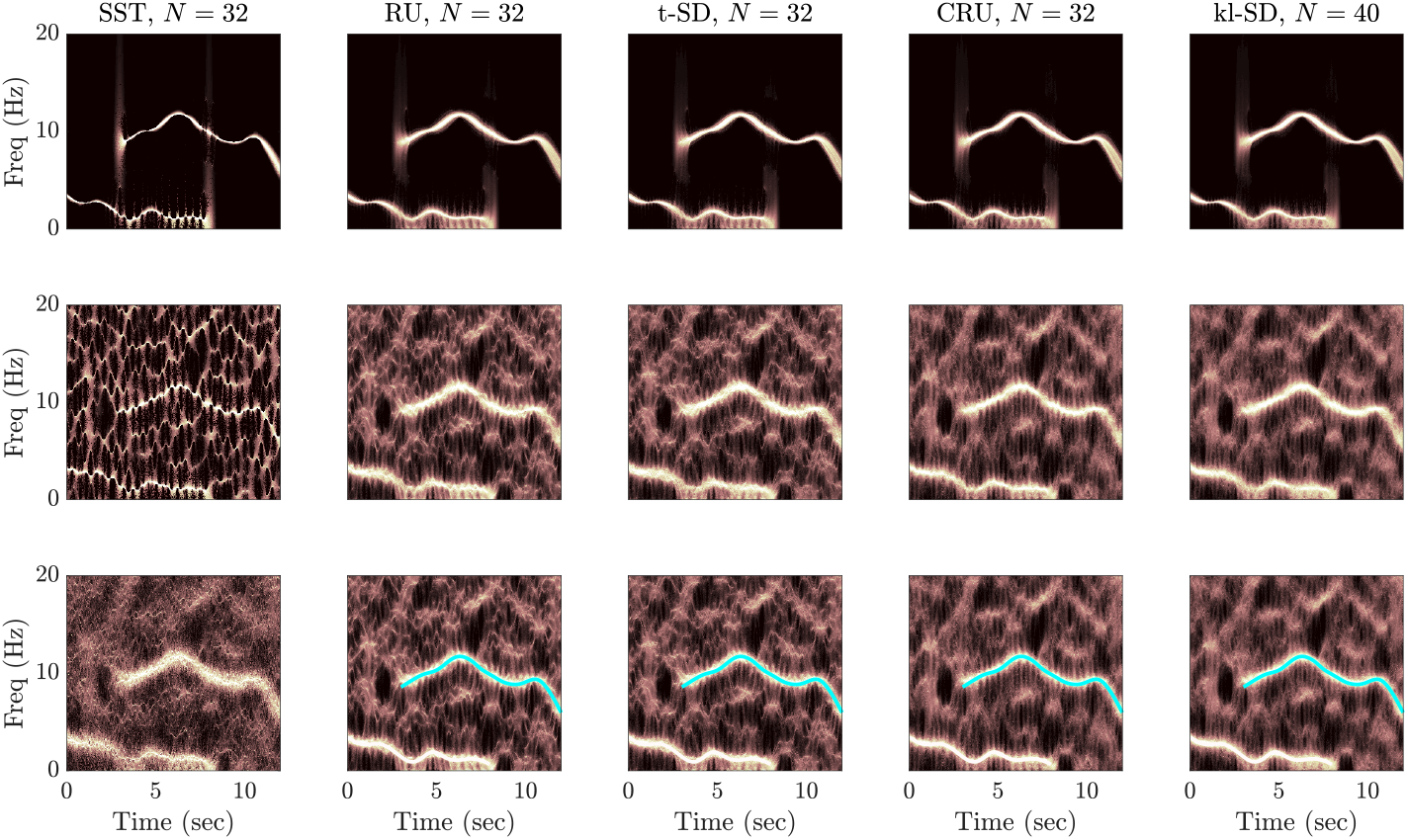
TFR’s by SST, MT-SST and various ConceFT’s for clean signal *f* (*t*) and noisy signal *Y*(*t*) over *t* ∈ [1/*ω*_0_, 16], where the sampling rate is *ω*_0_ = 100 (Hz) and the SNR is 0.11 dB. In the first column, from top to bottom shows the TFR’s of the clean signal *f* (*t*) analyzed by SST, the noisy signal *Y*(*t*) analyzed by SST, and the noisy signal *Y*(*t*) analyzed by MT-SST. From columns 2 to 5, we show TFR’s of the clean and noisy signals analyzed by RU-ConceFT, QU-ConceFT, the CRU-ConceFT, and CQU-ConceFT respectively. The first row is for the clean signal *f* (*t*), the second row is for the noisy signal *Y* (*t*), and the third row is for the noisy signal with the ground truth superimposed. For RU and CRU, we take 32 random points on 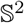 and Ω^3^. For QU-t-SD, we take the 7-design with 32 nodes on 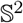. For QU-kl-SD, we take the triangle complex 4-design with 40 nodes on Ω^3^.

In Figure 2, it is observed that the information of interest, the two curves representing the instantaneous frequencies 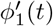 and 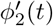, can be easily identified in the TFR of *f*(*t*) determined by SST. However, the TFR of *Y*(*t*) determined by SST is not suitable to identify the information of interest since the two curves representing instantaneous frequencies 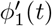 and 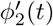 are buried in the background noise. The TFR determined by MT-SST provides more identifiable curves, but they are widened and blurred. This blurring is due to taking 6 Hermite windows, since the more the Hermite windows we take, the more spreading the Hermite windows is in the TF domain [3]. visually, the various Con-ceFT’s all enhance the contrast of the information of interest and the curves are sharpened. Specifically, both CRU-ConceFT and QU-ConceFT provide the sharpest curves. These results indicate that CQU-ConceFT performs almost equally compared with RU-ConceFT. in the next subsection, we provide a quantification of this visual finding to compare the performance of variations of ConceFT.

### 3.3. Performance measurement

To evaluate the performance of ConceFT’s with various designs, we follow the evaluation scheme proposed in [4, Section 4b], where the optimal-transport distance (OTD) between the *ideal TFR* of the clean signal and the TFR of the noisy signal is used to measure how accurate an algorithm can approximate the clean signal. The ideal TFR of *f* in (2) is defined as

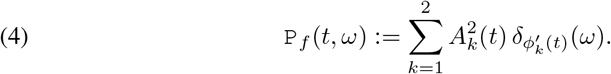

Let 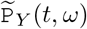 be the TFR determined by ConceFT for a stochastic signal. As 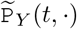 might not have integral 1 for time *t*, we normalize them such that 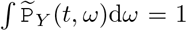. To simplify the notation, we use the same symbol for the normalized 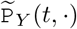. Then evaluate the OTD between 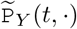 and `P`_*f*_ (*t*, ·), and average all the OTD’s over all sampling times to illustrate the quality of the estimator 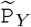 for `P`_*f*_.

Figure 3 shows the OTD for TFR’s of QU-ConceFT and CQU-ConceFT, and RU-ConceFT and CRU-ConceFT on 100 realizations of *Y* (*t*). Here we generate stochastic signal *Y* (*t*) for *t* ∈ [1/*ω*_0_,16] with the sampling rate *ω*_0_ = 100 (Hz) and the SNR is around 0.11 dB. As in Section 3.2, the QU-t-SD and RU are on 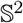 and QU-kl-SD and CRU are on Ω^3^. For RU-ConceFT and CRU-ConceFT, we realize up to 240 RU and CRU points on 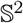 and Ω^3^ for 100 times and evaluate the mean and standard deviation of OTD’s, as indicated by the red and maroon in the figure. For QU-t-SD, we use up to the 21-design with nodes *N* ≤ 243. For QU-kl-SD, we use up to the 7-design with nodes *N* ≤ 254. We again use q =1 for all cases. In the real case, the mean and SD of the OTD’s for QU-ConceFT are both smaller than RU-ConceFT. in the complex case, CQU-ConceFT has a smaller mean than that of CRU-ConceFT while their standard deviations are almost the same. Moreover, the complex cases have a better performance than the real cases, which is consistent with the results of Section 3.2. These results support that the proposed (C)QU-ConceFT has a stable and better or at least equivalently good performance in approximating the ideal TFR of the clean signal compared with the RU-ConceFT with random points. We mention that the performance of different designs can be partially explained by the numerical integration approximation quality of the designs for the ideal ConceFT.

**Figure 3.**
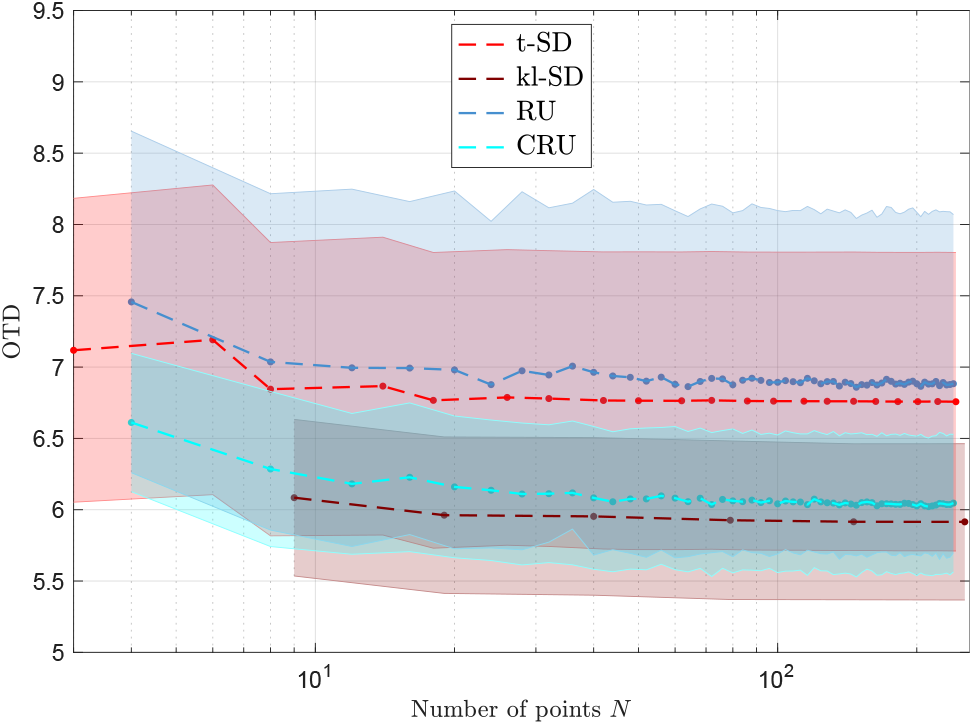
OTD’s of TFR’s of QU-ConceFT, CQU-ConceFT, RU-ConceFT and CRU-ConceFT for 100 stochastic signals *Y*(*t*) with the white Gaussian noise *ξ*(*t*) and *q* =1, where *t* ∈ [1/*ω*_0_,16], the sampling rate is *ω*_0_ = 100 (Hz) and the SNR is around 0.11 dB. The regions of light red, light maroon, light blue and light cyan show the variances for QU-t-SD, QU-kl-SD, RU, and CRU cases respectively. For the real cases, like QU-t-SD and RU, we take points on 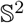. For the complex cases, like QU-kl-SD and CRU, we take points on Ω^3^.

Figure 4 shows the relation between the SNR and OTD for various QU-ConceFT’s. The realization of *Y* (*t*) is sampled at *ω*_0_ = 100 Hz from *t* ∈ [1/*ω*_0_,16]. The SNR ranges from −7 to 7. As above, the QU-t-SD uses the 7-design with 32 nodes on 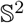 and the QU-kl-SD uses the triangle complex 4-design with 40 nodes on Ω^3^. On the other hand, the RU and CRU use 32 random points on 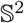 and Ω^3^ respectively. The behavior of the four cases are similar, that is, the OTD increases as the SNR increases. In the real case, QU-ConceFT and RU-ConceFT have almost equal OTD at each SNR value, while in the high SNR region, RU-ConceFT outperforms QU-ConceFT. In the complex case, CQU-ConceFT has a smaller OTD compared with CRU-ConceFT, while in the high SNR region, CRU-ConceFT outperforms CQU-ConceFT. In general, the real cases have higher OTD than the complex cases.

**Figure 4.**
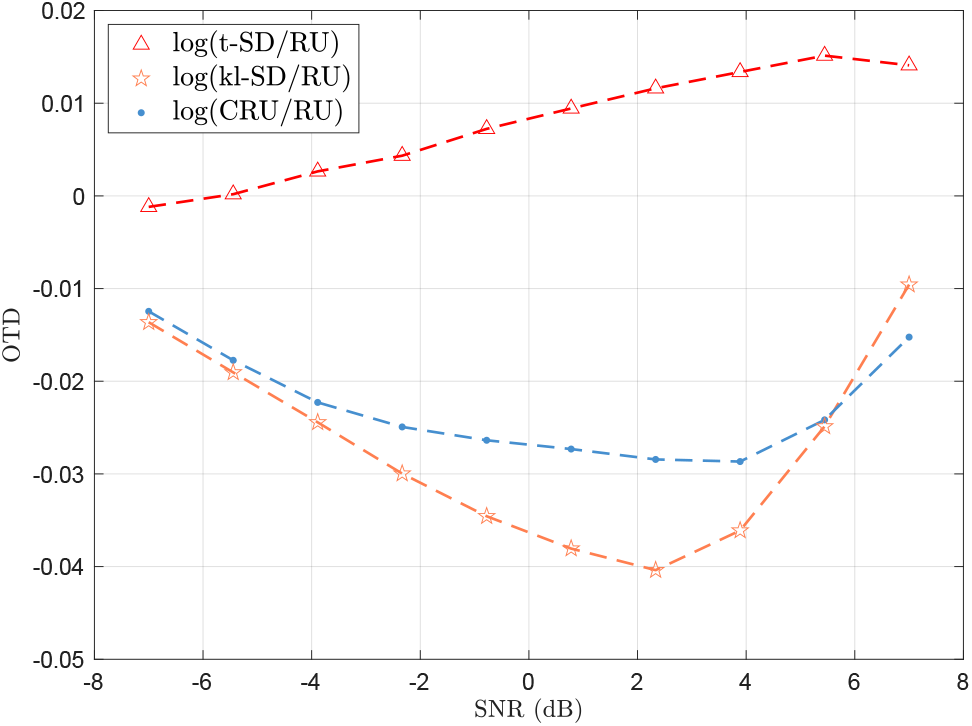
Illustration of the relationship between OTD and SNR for various TF analysis tools, including QU-ConceFT, CQU-ConceFT, RU-ConceFT and CRU-ConceFT. We take *q* = 1 in all ConceFT’s. in the plot, the y-axis is the ratio of OTD’s determined by different TF analysis tools divided by that determined by RU-ConceFT. The smaller the ratio OTD is, the more improvement is achieved. The stochastic signal *Y*(*t*) is sampled at ω_0_ = 100 Hz from *t* ∈ [1/*ω*_0_,16]. We take a 7-design with 32 nodes on 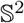 for the QU-t-SD and a triangle complex 4-designs with 40 nodes on Ω^3^ for the QU-kl-SD. For the RU and CRU, we take 32 randomly uniform points on 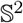 and Ω^3^ respectively.

## 4. Discussion and Conclusion

We introduce a novel generalized multitaper algorithm by combining a novel SD scheme and the nonlinear-type TF analysis that standardizes the RU-ConceFT. Its potential can be seen from an EEG signal analysis. For the practical purpose, we can apply CQU-ConceFT to study the EEG signal during the N2 sleep stage. One specific future mission is developing an automatic system to identify spindles based on CQU-ConceFT. The developed system will distinguish among different types of spindles by reading its TFR, and show how it is related to the sleep dynamics. We will report these clinical studies in future work.

## 5. Application to EEG signals

We apply QU-ConceFT to analyze the EEG signal during the N2 sleep stage and compare the results with SST and RU-ConceFT. We focus on the C3A2 EEG channel that was recorded at the sampling rate of 200Hz. The signal was recorded from a normal subject without sleep apnea by Alice 5 data acquisition system (philips Respironics, Murrysville, pA) in the sleep center at Chang Gung Memorial Hospital, Linkou, Taoyuan, Taiwan. The overnight sleep stages for all 30 seconds epochs, including Awake, REM, N1, N2, and N3, are provided by the consensus of two sleep experts. A typical feature of the EEG signal during the N2 stage is the spindle. Sleep spindles are brain activity bursts that oscillate at a frequency range of 11 to 16 Hz with a duration of 0.5 seconds or greater [5]. The nomination “spindle” comes from the fact that the “shape” of a sleep spindle is often like that of a yarn spindle. In Figure 5, we show a typical EEG epoch of 30 seconds long during the N2 sleep stage. The red arrows indicate the spindle events labeled by the sleep experts. We also show CQU-ConceFT, RU-ConceFT, and SST for the signal. It is clear that the labelled spindles oscillate at about 14 Hz, but with time-varying frequency. Moreover, three events, indicated by the blue arrows, look like a superposition of a spindle and a slow wave. This kind of pattern, in general, is *not* considered a spindle according to the American Academy of Sleep Medicine criteria [8]. However, the results well suggest that they are spindles; for example, see the dashed arrows in the second subplot. This “hidden” spindle structure is less discussed in the sleep literature, which might be due to the lack of proper analysis tools, while it might contain useful physiological information. However, a full exploration of this topic is out of the scope of this paper. Overall, CQU-ConceFT performs at least equally well compared with RU-ConceFT from the visualization perspective, while the windows are standardized via SD without any randomness. Moreover, compared with SST, both ConceFT’s have fewer “artifacts” indicated by the orange arrows.

**Figure 5.**
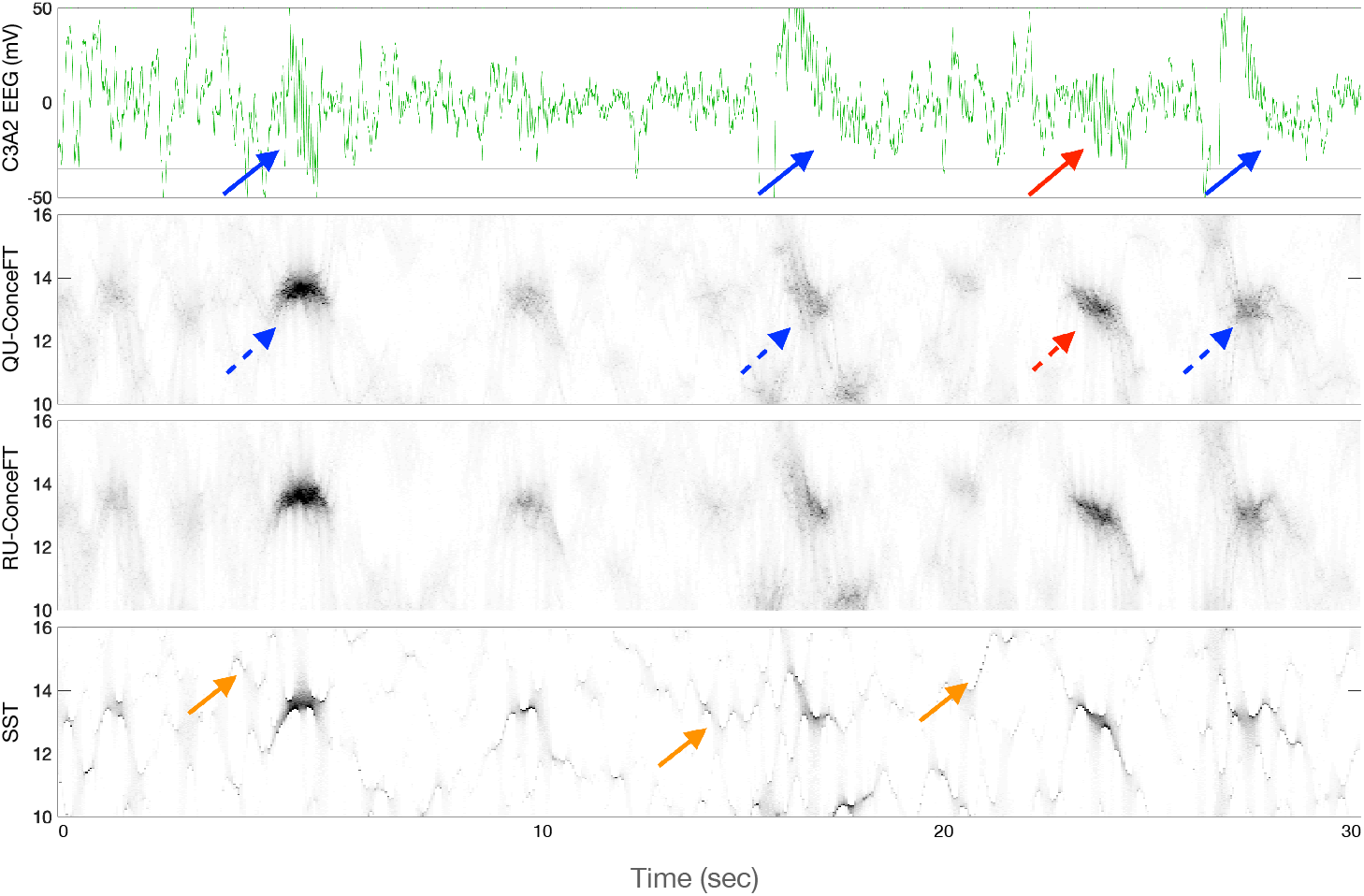
From top to bottom: an EEG epoch during the N2 stage, CQU-ConceFT, RU-ConceFT, and SST. We observe an annotated spindle around the 25th second, which we can visualize in all TFRs. However, it is visually clear that both ConceFT’s provide TFR’s with less “speckles” (indicated by orange arrows) compared with that generated by SST. Also, it is apparent that CQU-ConceFT performs at least equally well compared with RU-ConceFT.

## Footnotes

1 https://web.maths.unsw.edu.au/~rsw/Sphere/EffSphDes/

